# Functional coherence of the *Xist* and *RSX* protein interactomes: X chromosome inactivation in marsupials

**DOI:** 10.1101/2023.10.08.560959

**Authors:** Kim L. McIntyre, Shafagh A. Waters, Ling Zhong, Gene Hart-Smith, Mark Raftery, Jennifer A. Marshall Graves, Paul D. Waters

## Abstract

Long range epigenetic silencing is epitomised by X chromosome inactivation (XCI) in mammals. It is mediated by independently evolved, non-homologous long noncoding RNAs in eutherian and marsupial mammals (*XIST* and *RSX*). The *Xist* interactome, comprising proteins that mediate the silencing process, is well documented in mouse studies. Here we interrogate proteins that interact with *RSX* using chromatin isolation by RNA purification followed by mass spectrometry (ChIRP-MS) in a marsupial representative, *Monodelphis* domestica. We identify 135 proteins that interact with *RSX*, of which 56 have orthologues in the *Xist* interactome. Remarkably, nearly 90% of the combined *Xist* and *RSX* interactomes are within the same protein-protein association network. This network clustered into three major groups with distinctive functional enrichments, including RNA splicing and processing, regulation of translation and ribosomal biogenesis, and epigenetic transcriptional silencing. The proteins of the *RSX* interactome were enriched for regions of intrinsic disorder in common with the *Xist* interactome, identifying this as a feature of ribonucleoprotein complexes associated with XCI. We also show that RNAi knockdown of representative *RSX* interactors, HNRNPK and CKAP4, led to reactivation of transcription from the inactive X chromosome, indicating a role for each in marsupial XCI. Thus, despite the absence of linear sequence homology between *Xist* and *RSX*, they exhibit extraordinary functional coherence that indicates potential for post-transcriptional regulation, a feature not previously associated with the molecular machinery of XCI.

## Introduction

The sex chromosomes of therian mammals (marsupials and eutherians) share a common ancestry (*1*), having evolved from a pair of autosomes (*2*) after the divergence of therian and monotreme mammals approximately 187 mya (*3*). X chromosome inactivation (XCI) occurs in both groups of mammals, implying an ancient origin. However, XCI in eutherians and marsupials involve molecular mechanisms that are remarkably different.

XCI in therian mammals silences transcription of one of the two X chromosomes in female somatic cells (*4*). It is established in the early embryo and maintained through subsequent cell divisions, and serves as an important model for epigenetic silencing due to its unparalleled scale and stability. Long-noncoding RNAs (lncRNAs) have emerged as common regulators in therian XCI. Eutherian XCI is mediated by a lncRNA called *XIST* (*5*). Its mouse ortholog, *Xist* (*6*), shares &#x223C;67% sequence conservation with human *XIST*. This includes a series of tandem repeats (A to F), of which only repeat A is well conserved across all eutheria (*7*).

The protein interactome of *Xist* has been investigated in mouse cell lines using techniques involving chromatin isolation by RNA precipitation with mass spectrometry (ChIRP-MS) and its variations (*8–12*). These investigations have identified 494 proteins in total, with only 6 proteins (Hnrnpm, Hnrnpu, Myef2, Raly, RBM15, Spen) common to all studies (*8–10, 12*). An alternative technique, RNA immunoprecipitation (RIP) combined with deep sequencing, identified epigenetic regulators in the human *XIST* interactome that were not identified in the mouse studies: EZH2 and SUZ12, subunits of polycomb repressive complex 2 (PRC2), and CHD4, a subunit of the NuRD histone deacetylase complex (*13, 14*).

*Mars*upials lack an *XIST* gene; instead, ancient protein coding genes have been retained at the loci homologous to those from which *XIST* and neighbouring genes evolved in eutherians (*15–17*). In marsupials, XCI is mediated by a lncRNA called *RSX* (*18*) that derives from a non-homologous and physically distinct region of the X chromosome. *RSX* is 27 kb in *Monodelphis domestica* (*18*) and 30 kb in koala (*Phascolarctus cinereus*) (*19*), longer than the 15 kb mouse *Xist* (*6*) and the 17 kb human *XIST* (*20*). Although lacking linear sequence homology, a k-mer analysis classified two major groupings of repeat domains that are shared between *Xist* and *RSX* (*RSX* repeat 1 with *Xist* repeats B, C and *XIST* repeat D, and *RSX* repeats 2, 3 and 4 with *Xist* repeats A and E). Each of these domains are enriched for specific protein binding motifs (*21*). Therefore, although *RSX* and *Xist* share no sequence homology they could be functional analogs.

*Xist* and *RSX* are both nuclear transcripts that are spliced, capped and polyadenylated in the manner of mRNAs, and are expressed only in female somatic cells, exclusively from the inactive X chromosome. In both cases, the clustered transcripts can be visualised using RNA fluorescence in-situ hybridization (RNA FISH) as a distinctive cloud-like signal accumulated on the inactive X chromosome (*18, 20*). Induction of *RSX* expression from an autosomal transgene in mouse silences transcription *in cis* (*18*). This indicates a silencing capacity similar to *Xist* (*22*), although marsupial XCI is ‘leakier’ or more incomplete than the *XIST*-driven process in eutherians (*23*), perhaps due to the evolution of two different lncRNAs in different ancestral genomic contexts.

Here, we investigate the protein interactome of *RSX* in a marsupial, *M. domestica*, and compare it with the *Xist* protein interactome. We consider the molecular mechanisms underlying the convergent evolution of XCI in therian mammals to enhance understanding of the evolution of the adaptations for balancing gene expression between the sexes. Our findings show that *RSX* interactors significantly overlap with *Xist* interactors, falling within the same protein-protein association network related to RNA splicing and processing, translation regulation and ribosome biogenesis, and epigenetic transcriptional silencing. This highlights the remarkable functional coherence of these non-homologous and independently evolved lncRNAs. We identified overlap between the *Xist* and *RSX* protein interactomes, both of which are enriched for functions associated with post-transcriptional regulation of gene expression. Post-transcriptional regulation has been shown to contribute to the balancing of expression of X-borne genes between the sexes in eutherians (*24–27*), although the underlying mechanisms are unknown.

## Results

### Identification and validation of *RSX* interactors and comparision with *Xist* interactome

To investigate the protein interactome of *RSX* we used ChIRP-MS to capture proteins associated with *RSX* using six biotinylated oligonucleotides complementary to different *RSX* regions. Cell lysates were prepared from female *M. domestica* fibroblast cells that were either UV crosslinked, formaldehyde crosslinked, or uncrosslinked. We identified 131 proteins that associated with *RSX* using alternate criteria of presence/absence and greater than two-fold enrichment versus a control, either absence of oligonucleotides or scrambled oligonucleotides (Figure 1A).

**Figure 1.**
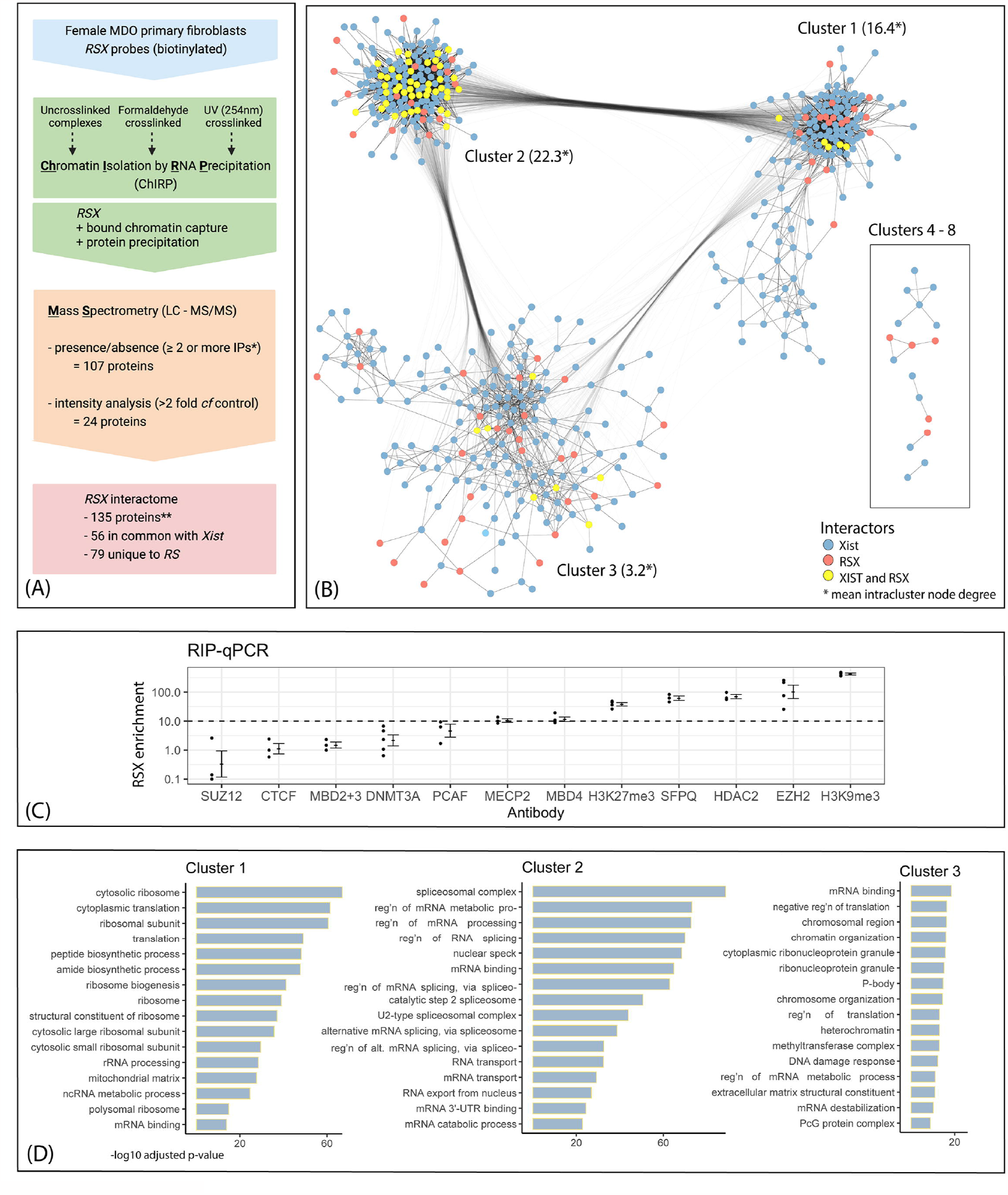
*RSX* and *Xist* protein interactomes share common orthologous proteins and protein-protein association networks with distinctive functional enrichments. (**A)** Overview of ChIRP-MS workflow. *Two proteins were identified by a single pulldown from a UV crosslinked sample. **includes 4 additional proteins identified using RIP-qPCR. **(B)** Protein-protein interactions of the *RSX* and *Xist* interactomes based on experimentally determined interactions, co-expression, and curated database annotations for human orthologs (STRING database v11.5) (*29*). Each node represents an interactome protein, each edge represents an annotated protein-protein interaction of minimum confidence 0.4. Interaction networks were visualised using Cytoscape (v3.8.2), omitting proteins with no annotated interactions. Nodes were clustered based on connectivity (number and weight of edges) using the GLay Cytoscape plugin (*42*) with default settings (prefuse force directed layout). Intercluster edges to minor clusters (4-10) omitted for clarity. (**C)** Enrichment of *RSX* (fold change relative to Igg controls) by immunoprecipitation of protein targets from female *M. domestica* fibroblast cell lysates, followed by quantitative PCR using *RSX*-specific primers. Enrichment (30-fold) was also detected for HNRNPK, as previously published (*21*). (**D**) *Key* functional and structural enrichments of each major protein interaction cluster. GSEA was conducted using gProfiler2 (*3*) with multiple testing correction based on false discovery rate.

We validated two *RSX* interactors using RNA immunoprecipitation (RIP) followed by quantitative PCR (qPCR). We also assayed *RSX* interaction with eight known chromatin modifiers and two repressive histone marks (Figure 1C). Target proteins were immunoprecipitated from female *M. domestica* fibroblast cell lysates, followed by qPCR using *RSX*-specific primers (*21*). Enrichment of *RSX* (relative to Igg controls) of greater than 10-fold was detected for seven targets (Figure 1C). These included HNRNPK (*21*) and SFPQ, which were identified by the ChIRP-MS. The RIP-qPCR also identified four additional *RSX* interactors not detected by ChIRP-MS: EZH2 (PRC2 catalytic subunit), HDAC2 (a histone deacetylase), MBD4 (a methyl-CpG binding domain protein), and MECP2 (a methyl-CpG-binding protein).

These proteins were included in the *RSX* interactome, bringing the total to 135. Of these, 79 were specific to the *RSX* interactome. However, the other 56 had orthologs that were identified in the *Xist* interactome (497 proteins), and 33 *RSX*-interactors had orthologs that were identified in at least two of the *Xist* interactome studies. Therefore, we identified a substantial cohort of proteins that interact with both *RSX* and *Xist*, despite the lack of homology in primary sequence of these two lncRNAs. We also considered the extent to which the two interactomes might include different proteins from common functional pathways, potentially providing insights to understanding how therian XCI evolved to be mediated by different lncRNAs in marsupials and eutherians.

### Network analysis of the *RSX* and *Xist* interactomes reveals functional similarities

Gene set enrichment analysis (GSEA) (*28*) of each of the *RSX* and *Xist* interactomes identified that nearly 90% of the 135 ontology terms enriched for the *RSX* interactome were also enriched for the *Xist* interactome (p < 1 x 10^-3^), suggesting functional similarities between the two interactomes.

We queried the protein-protein interactions within the combined *RSX* and *Xist* interactomes using the STRING database (v11.5) (*29*). Of the 576 proteins in the combined interactomes, 516 proteins had at least one interaction (confidence score > 0.4) and formed a network with 8,701 edges (a mean of 15.1 edges per node). This was significantly higher than the 3,599 edges expected for a random set of 576 proteins (p < 1 x 10^-16^). Clustering of the interaction network partitioned it into three larger clusters and five small clusters. The key functional enrichments of each of the three major clusters were determined using GSEA. The three large clusters were individually enriched for functions including mRNA binding, translation (and regulation of translation), and nitrogen compound catabolic process (Figure 1D). In addition to these common terms, the clusters had distinctive functional enrichments, including ribosomal biogenesis in cluster 1, RNA splicing and processing in cluster 2, and chromatin modification and epigenetic silencing in cluster 3.

Clustering and enrichment analyses were also conducted on the *RSX* and *Xist* interactomes separately using the same approach. Each interactome had four major clusters, with GSEA enrichments reflecting those of the combined interactomes analysis, subject to division of cluster 2 in the *RSX* interactome, and division of cluster 1 in the *Xist* interactome.

*RSX*-specific interactome proteins were of interest in unravelling the differences between eutherian and marsupial XCI. GSEA of the 79 *RSX*-specific proteins identified enrichments for spliceosomal complexes, ribosomal subunits, cytosolic translation, nucleosome binding and chromatin organization in proportions similar to those of the overall *RSX* interactome, other than perhaps for nucleosome binding which predominantly involves *RSX*-specific proteins. Apart from this, *RSX*-specific proteins did not appear to have gross unique function compared to the full *RSX* interactome.

Collectively, the clustering and GSEA enrichment analyses revealed an overlap between the *RSX* and *Xist* interactomes. This encompassed common proteins and also interactions with different proteins in common molecular pathways, providing insights into the functions modulated by *RSX* and *Xist*.

### Functional analysis of HNRNPK and CKAP4 in marsupial XCI

We focused on the functional role of HNRNPK, which was identified in our *RSX* interactome and is also an *Xist*-interacting protein. HNRNPK is important in recruiting polycomb repressive complex 1 (PRC1), a significant part of the epigenetic silencing machinery, during eutherian XCI (*11, 30*). In the combined *RSX/Xist* interactome network it was in cluster 2, which was enriched for functions in RNA splicing and processing (Figure 1B). We depleted HNRNPK expression in a female *M. domestica* fibroblast cell line using RNA interference (RNAi) and assessed the effect on XCI using RNA FISH. This allowed us to determine transcriptional status of *MSN*, an X-borne gene that is usually silenced on the inactive X chromosome, so should only have monoallelic expression. In wild-type control nuclei (transfected with an empty RNAi vector) biallelic expression of *MSN* (indicating transcription from both X chromosomes) was detected in only 18% of cells (n = 286; Figure 2A).

**Fig. 2.**
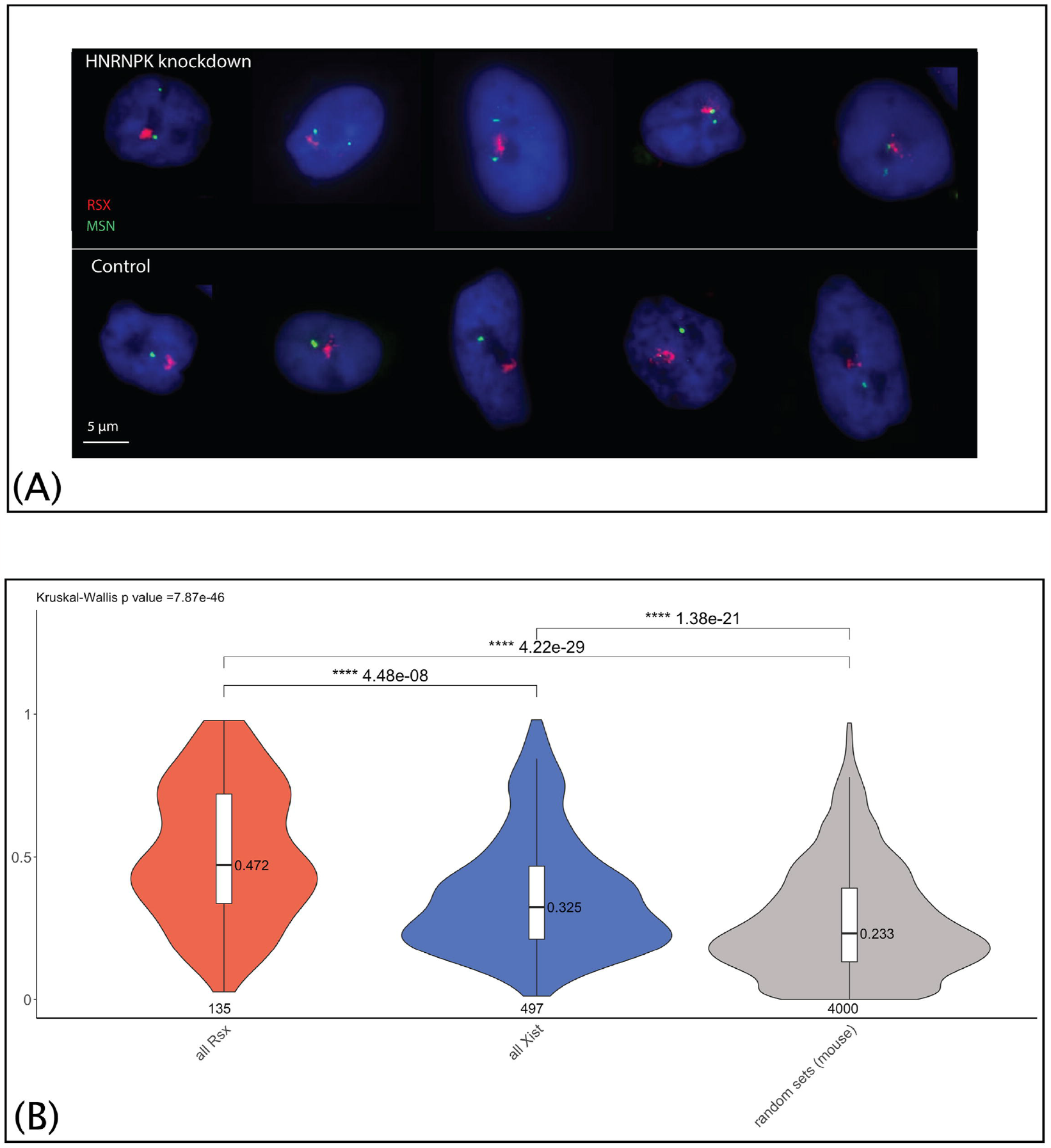
*RSX*-interacting protein, HNRNPK, has a role in maintaining XCI. The *RSX* and *Xist* interactomes are both enriched for proteins with intrinsically disordered regions. **(A)** RNAi knockdown of HNRNPK (∼24-35% knockdown efficiency) led to reactivation of transcription from the inactive X chromosome, evident as two *MSN* signals (p = 2.7 x 10^-6^ (Fisher’s Exact Test), p = 1.0 x 10^-11^ (Chi Squared Test Goodness of Fit Test), n = 159). Dual color RNA FISH using probes for *RSX* (red), and the X-borne gene, *MSN* (green) in female *M. domestica* fibroblasts, selected images. (**B)** Median protein IDR scores for the *RSX* and *Xist* interactomes represented as violin plots (depicting density distribution) overlayed with boxplots depicting the median for all proteins of the *RSX* and *Xist* interactomes (mouse orthologs), and randomly sampled proteins of the mouse proteome (20x sets of 200 proteins combined). Statistical significance assessed using Dunn’s Test (with Holm adjustment) for pairwise comparisons, following Kruskal-Wallis Test.

Following HNRNPK knockdown (by ∼24-35%) biallelic expression of *MSN* increased from 18% in control cells to 39% in cells with depleted HNRNPK expression (n = 159; p = 1.0 x 10^-11^ Chi Squared Test Goodness of Fit Test; p = 2.7 x 10^-6^ Fisher’s Exact Test) (Figure 2A). Increased biallelic expression of *MSN* signified reactivation of transcription from the silenced allele on the inactive X chromosome. This outcome was consistently observed across two independent experiments.

We also evaluated CKAP4 which, unlike HNRNPK, has not been identified as an *Xist* interactor, and has no predicted interactions with any protein in either the *Xist* or *RSX* interactomes. In our *RSX* interactome, CKAP4 was the protein with the highest fold-change (log2) enrichment relative to each of the controls in the native (uncrosslinked) ChIRP-MS. We used RNAi to suppress CKAP4 in female *M. domestica* fibroblasts by ∼53-55%. We observed an increase in biallelic expression of *MSN* from 18% to 26% (n = 165; p = 6.0 x 10^-2^ Fisher’s Exact Test; p = 8.7 x 10^-3^ Chi Squared Test Goodness of Fit Test). This suggests that CKAP4 plays a role in marsupial XCI.

Interestingly, despite the absence of CKAP4 from the GSEA analysis of the combined interactome network, the rough endoplasmic reticulum (where CKAP4 usually localises) was significantly enriched in cluster 1 (p = 9.8 x 10^-6^), along with three other associated terms (p < 8.0 x 10^-4^). This aligns with the functional enrichment of ribosomal and translation-associated machinery observed in the same cluster.

These data provide evidence that both HNRNPK and CKAP4 play roles in transcriptional silencing of the inactive X chromosome in marsupials. Characterising other *RSX* interactors will help us to fully understand the molecular mechanisms underlying their function in X chromosome inactivation.

### Enrichment of proteins with Intrinsically Disordered Regions (IDRs) in the *RSX* interactome

Recent studies have revealed that the *Xist* compartment is founded on an assembly of dynamic RNP complexes comprising *Xist* RNA in association with the intrinsically disordered regions (IDRs) of *Xist*-interacting proteins, such as SPEN, PTBP1, MATR3, CELF1, CIZ1 (*31–34*). Of these, only PTBP1 was identified in the *RSX* interactome, so we considered whether other proteins with IDRs might also be present in the *RSX* interactome.

We assessed the proportion of proteins enriched for IDRs in each interactome with IUPred2A (*35*), which calculates a disorder score for each residue using amino acid composition and energy estimation. Disorder scores above 0.5 (range 0 to 1) correspond to disordered residues. We calculated the IDR score for each protein as the median disorder score of its residues. We found that *RSX* interactome proteins had higher median IDR scores than the *Xist* interactome proteins (p = 4.5 x10^-8^). Both interactomes had higher median IDR scores than a randomly sampled group of 200 proteins from the reference proteome (p = 4.2 x10^-29^ *cf RSX* and p = 1.4 x10^-21^ *cf Xist*) (Figure 2B). Proteins common to both the *RSX* and *Xist* interactomes had higher median IDR scores than proteins that interact exclusively with either *RSX* or *Xist* (p = 2.1 x10^-2^ *cf RSX*, p = 5.5 x10^-10^ *cf Xist*).

IDRs have important roles supporting protein-protein interactions, protein-RNA interactions, and the formation of phase-separated condensates that form nuclear subcompartments (*36–38*). Our finding suggests that an enrichment for proteins with IDRs may play a role in the formation of *RSX*-associated RNP complexes, aiding subcellular organisation, as has been identified for *Xist*-associated RNPs.

## Discussion

This research provides a novel insight into the complex protein interactions of *RSX*, a lncRNA in marsupials with a role similar to the eutherian *Xist* in epigenetically silencing the inactive X chromosome. We found that that the *RSX* interactome has functional enrichments analogous to *Xist* that underscore their functional similarities. We also showed that alleles on the inactive X chromosome were partially reactivated following the partial depletion of HNRNPK and CKAP4, two proteins in the *RSX* interactome, indicating a role for each in marsupial XCI.

LncRNAs provide an important organizing mechanism in epigenetic regulation, including in the recruitment and sequestration of RNA splicing and processing factors. Many of these proteins are multifunctional, often with distinct nuclear and cytoplasmic functions. This interaction of lncRNAs with multifunctional proteins can provide an efficient mechanism by which lncRNAs can impact diverse molecular networks. Of the 56 proteins identified in common in the *RSX* and *Xist* interactomes, 44 form part of interactome network cluster 2, which features proteins involved in RNA splicing and processing.

The 56 proteins common to both interactomes were enriched for intrinsically disordered regions (IDRs), a characteristic identified in each interactome individually. These IDRs could potentially contribute to epigenetic silencing by facilitating protein-protein interactions, as proteins enriched in IDRs are characterised by their flexible and adaptable binding with multiple partners. This binding plasticity may contribute to the dynamic regulation of gene expression on the inactive X chromosome, potentially including alternate silencing and escape from silencing, depending on specific cellular contexts. Further, this plasticity might be important for the observed ‘leakiness’ of XCI observed in marsupials as partial or full expression from the inactive X chromosome (*39, 40*).

Analysis of interactome network clusters 1 and 3 provided further insight into the mechanisms by which *RSX* and *Xist* might regulate gene expression. GSEA of each of these clusters indicated a functional coherence: despite a relatively small overlap in interactomes, they had different protein interactors involved in shared pathways. Cluster 3 contained proteins typically associated with XCI, including those involved in epigenetic regulation of transcriptional silencing, histone modifications and heterochromatin. Cluster 1 was functionally enriched for ribosomal biogenesis, rRNA processing and regulation of translation. For *RSX*, this is consistent with the nucleolar association of the inactive X in marsupials.

Enrichment of functions associated with post-transcriptional regulation is another fascinating aspect of the *Xist* and *RSX* interactomes. Post-transcriptional regulation of X-borne gene expression has been identified in eutherians and *M. domestica* by comparing gene expression in the transcriptome and translatome (based on ribosomal occupancy) (*24*). Balancing of sex chromosome-borne gene expression between sexes in the proteome has also been identified in the more distantly related platypus (*Ornithorhynchus anatinus*) and chicken (*Gallus gallus*). This highlights the possibility of post-transcriptional regulation being an ancestral strategy for fine tuning expression of sex chromosome genes in both sexes (*41*).

*XIST* and *RSX* provide a striking example of convergent evolution: independently evolved lncRNAs that recruit similar molecular pathways to repress the activity of almost an entire chromosome. Our understanding of the evolution of the therian sex chromosomes suggests that silencing of X-borne genes would have evolved as the X and Y chromosomes diverged, perhaps initially involving other noncoding RNAs and only localised silencing before the Y chromosome was as degraded as it currently is. The emergence of independent chromosome wide regulation of XCI by *XIST* and *RSX* would then have coordinated presumably more efficient silencing, balancing the expression of X-borne genes between the sexes as degeneration of the Y chromosome progressed.

## Acknowledgements

Not applicable.

## Author Contributions

K.L.M. drafted the manuscript and figures, performed RNA FISH, and conceived and performed IDR and GSEA analysis. S.A.W. conceived and performed ChIRP and RIP-qPCR, and performed manuscript editing and figure preparation. L.Z. generated mass spectrometry data. G.H.S. performed mass spectrometry data analysis. M.R. was involved in mass spectrometry data collection and analysis. J.A.M. commented on and edited the manuscript. P.D.W. conceived and oversaw the project and the assembly of the manuscript.

## References

1. J. A. M. Graves, The evolution of mammalian sex chromosomes and the origin of sex determining genes. Philos Trans R Soc Lond B Biol Sci. 350 (1995), doi:10.1098/rstb.1995.0166.

2. S. Ohno, Sex chromosome and sex-linked genes (Springer-Verlag, Berlin, Heidelberg, Berlin, 1967).

3. Y. Zhou, L. Shearwin-Whyatt, J. Li, Z. Song, T. Hayakawa, D. Stevens, J. C. Fenelon, E. Peel, Y. Cheng, F. Pajpach, N. Bradley, H. Suzuki, M. Nikaido, J. Damas, T. Daish, T. Perry, Z. Zhu, Y. Geng, A. Rhie, Y. Sims, J. Wood, B. Haase, J. Mountcastle, O. Fedrigo, Q. Li, H. Yang, J. Wang, S. D. Johnston, A. M. Phillippy, K. Howe, E. D. Jarvis, O. A. Ryder, H. Kaessmann, P. Donnelly, J. Korlach, H. A. Lewin, J. Graves, K. Belov, M. B. Renfree, F. Grutzner, Q. Zhou, G. Zhang, Platypus and echidna genomes reveal mammalian biology and evolution. Nature. 592, 756–762 (2021).

4. J. A. M. Graves, S. M. Gartler, Mammalian X chromosome inactivation: Testing the hypothesis of transcriptional control. Somat Cell Mol Genet. 12, 275–280 (1986).

5. C. J. Brown, A. Ballabio, J. L. Rupert, R. G. Lafreniere, M. Grompe, R. Tonlorenzi, H. F. Willard, A gene from the region of the human X inactivation centre is expressed exclusively from the inactive X chromosome. Nature. 349, 38–44 (1991).

6. N. Brockdorff, A. Ashworth, G. F. Kay, V. M. McCabe, D. P. Norris, P. J. Cooper, S. Swift, S. Rastan, The product of the mouse Xist gene is a 15 kb inactive X-specific transcript containing no conserved ORF and located in the nucleus. Cell. 71, 515–526 (1992).

7. T. Dixon-McDougall, C. J. Brown, Independent domains for recruitment of PRC1 and PRC2 by human XIST. PLoS Genet. 17, 1–28 (2021).

8. C. Chu, Q. C. Zhang, S. T. da Rocha, R. A. Flynn, M. Bharadwaj, J. M. Calabrese, T. Magnuson, E. Heard, H. Y. Chang, Systematic discovery of Xist RNA binding proteins. Cell. 161, 404–416 (2015).

9. A. Minajigi, J. E. Froberg, C. Wei, H. Sunwoo, B. Kesner, D. Colognori, D. Lessing, B. Payer, M. Boukhali, W. Haas, J. T. Lee, A comprehensive Xist interactome reveals cohesin repulsion and an RNA-directed chromosome conformation. Science (1979). 349 (2015), doi:10.1126/science.aab2276.

10. C. A. McHugh, C. K. Chen, A. Chow, C. F. Surka, C. Tran, P. McDonel, A. Pandya-Jones, M. Blanco, C. Burghard, A. Moradian, M. J. Sweredoski, A. A. Shishkin, J. Su, E. S. Lander, S. Hess, K. Plath, M. Guttman, The Xist lncRNA interacts directly with SHARP to silence transcription through HDAC3. Nature. 521, 232–236 (2015).

11. G. Pintacuda, G. Wei, C. Roustan, B. A. Kirmizitas, N. Solcan, A. Cerase, A. Castello, S. Mohammed, B. Moindrot, T. B. Nesterova, N. Brockdorff, hnRNPK Recruits PCGF3/5-PRC1 to the Xist RNA B-Repeat to Establish Polycomb-Mediated Chromosomal Silencing. Mol Cell. 68, 955–969.e10 (2017).

12. A. Bousard, A. C. Raposo, J. J. Żylicz, C. Picard, V. B. Pires, Y. Qi, C. Gil, L. Syx, H. Y. Chang, E. Heard, S. T. da Rocha, The role of Xist Cmediated Polycomb recruitment in the initiation of XCchromosome inactivation . EMBO Rep. 20, 1–18 (2019).

13. D. G. Hendrickson, D. R. Kelley, D. Tenen, B. Bernstein, J. L. Rinn, Widespread RNA binding by chromatin-associated proteins. Genome Biol. 17, 28 (2016).

14. Z. Lu, J. K. Guo, Y. Wei, D. R. Dou, B. Zarnegar, Q. Ma, R. Li, Y. Zhao, F. Liu, H. Choudhry, P. A. Khavari, H. Y. Chang, Structural modularity of the XIST ribonucleoprotein complex. Nat Commun. 11, 1–14 (2020).

15. L. Duret, C. Chureau, S. Samain, J. Weissanbach, P. Avner, The Xist RNA gene evolved in eutherians by pseudogenization of a protein-coding gene. Science (1979). 312, 1653–1655 (2006).

16. T. A. Hore, E. Koina, M. J. Wakefield, J. A. M. Marshall Graves, The region homologous to the X-chromosome inactivation centre has been disrupted in marsupial and monotreme mammals. Chromosome Res. 15, 147–161 (2007).

17. A. I. Shevchenko, I. S. Zakharova, E. A. Elisaphenko, N. N. Kolesnikov, S. Whitehead, C. Bird, M. Ross, J. R. Weidman, R. L. Jirtle, T. V. Karamysheva, N. B. Rubtsov, J. L. VandeBerg, N. A. Mazurok, T. B. Nesterova, N. Brockdorff, S. M. Zakian, Genes flanking Xist in mouse and human are separated on the X chromosome in American marsupials. Chromosome Research. 15, 127–136 (2007).

18. J. Grant, S. K. Mahadevaiah, P. Khil, M. N. Sangrithi, H. Royo, J. Duckworth, J. R. McCarrey, J. L. VandeBerg, M. B. Renfree, W. Taylor, G. Elgar, R. D. Camerini-Otero, M. J. Gilchrist, J. M. Turner, Rsx is a metatherian RNA with Xist-like properties in X-chromosome inactivation. Nature. 487, 254–258 (2012).

19. R. N. Johnson, D. O'Meally, Z. Chen, G. J. Etherington, S. Y. W. Ho, W. J. Nash, C. E. Grueber, Y. Cheng, C. M. Whittington, S. Dennison, E. Peel, W. Haerty, R. J. O'Neill, D. Colgan, T. L. Russell, D. E. Alquezar-Planas, V. Attenbrow, J. G. Bragg, P. A. Brandies, A. Y. Y. Chong, J. E. Deakin, F. Di Palma, Z. Duda, M. D. B. Eldridge, K. M. Ewart, C. J. Hogg, G. J. Frankham, A. Georges, A. K. Gillett, M. Govendir, A. D. Greenwood, T. Hayakawa, K. M. Helgen, M. Hobbs, C. E. Holleley, T. N. Heider, E. A. Jones, A. King, D. Madden, J. A. M. Graves, K. M. Morris, L. E. Neaves, H. R. Patel, A. Polkinghorne, M. B. Renfree, C. Robin, R. Salinas, K. Tsangaras, P. D. Waters, S. A. Waters, B. Wright, M. R. Wilkins, P. Timms, K. Belov, Adaptation and conservation insights from the koala genome. Nat Genet. 50, 1102–1111 (2018).

20. C. J. Brown, B. D. Hendrich, J. L. Rupert, R. G. Lafrenière, Y. Xing, J. Lawrence, H. F. Willard, The human XIST gene: Analysis of a 17 kb inactive X-specific RNA that contains conserved repeats and is highly localized within the nucleus. Cell. 71, 527–542 (1992).

21. D. Sprague, S. A. Waters, J. M. Kirk, J. R. Wang, P. B. Samollow, P. D. Waters, J. M. Calabrese, Nonlinear sequence similarity between the Xist and Rsx long noncoding RNAs suggests shared functions of tandem repeat domains. Rna. 25, 1004–1019 (2019).

22. I. Naciri, B. Lin, C. H. Webb, S. Jiang, S. Carmona, W. Liu, A. Mortazavi, S. Sun, Linking Chromosomal Silencing With Xist Expression From Autosomal Integrated Transgenes. Front Cell Dev Biol. 9, 1–12 (2021).

23. S. Al Nadaf, J. E. Deakin, C. Gilbert, T. J. Robinson, J. A. M. Graves, P. D. Waters, A cross-species comparison of escape from X inactivation in Eutheria: implications for evolution of X chromosome inactivation. Chromosoma. 121, 71–78 (2012).

24. Z. Y. Wang, E. Leushkin, A. Liechti, S. Ovchinnikova, K. Mößinger, T. Brüning, C. Rummel, F. Grützner, M. Cardoso-Moreira, P. Janich, D. Gatfield, B. Diagouraga, B. de Massy, M. E. Gill, A. H. F. M. Peters, S. Anders, H. Kaessmann, Transcriptome and translatome coevolution in mammals. Nature. 588, 642–647 (2020).

25. S. Yin, W. Deng, H. Zheng, Z. Zhang, L. Hu, X. Kong, Evidence that the nonsense-mediated mRNA decay pathway participates in X chromosome dosage compensation in mammals. Biochem Biophys Res Commun. 383, 378–382 (2009).

26. A. J. Brenes, H. Yoshikawa, D. Bensaddek, B. Mirauta, D. Seaton, J. L. Hukelmann, H. Jiang, O. Stegle, A. I. Lamond, Erosion of human X chromosome inactivation causes major remodeling of the iPSC proteome. Cell Rep. 35, 109032 (2021).

27. M. L. Faucillion, J. Larsson, Increased expression of X-linked genes in mammals is associated with a higher stability of transcripts and an increased ribosome density. Genome Biol Evol. 7, 1039–1052 (2015).

28. U. Raudvere, L. Kolberg, I. Kuzmin, T. Arak, P. Adler, H. Peterson, J. Vilo, G:Profiler: A web server for functional enrichment analysis and conversions of gene lists (2019 update). Nucleic Acids Res. 47, W191–W198 (2019).

29. D. Szklarczyk, A. L. Gable, D. Lyon, A. Junge, S. Wyder, J. Huerta-Cepas, M. Simonovic, N. T. Doncheva, J. H. Morris, P. Bork, L. J. Jensen, C. Von Mering, STRING v11: Protein-protein association networks with increased coverage, supporting functional discovery in genome-wide experimental datasets. Nucleic Acids Res. 47, D607–D613 (2019).

30. D. Colognori, H. Sunwoo, A. J. Kriz, C. Y. Wang, J. T. Lee, Xist Deletional Analysis Reveals an Interdependency between Xist RNA and Polycomb Complexes for Spreading along the Inactive X. Mol Cell. 74, 101–117.e10 (2019).

31. Y. Markaki, J. Gan Chong, Y. Wang, E. C. Jacobson, C. Luong, S. Y. X. Tan, J. W. Jachowicz, M. Strehle, D. Maestrini, A. K. Banerjee, B. A. Mistry, I. Dror, F. Dossin, J. Schöneberg, E. Heard, M. Guttman, T. Chou, K. Plath, Xist nucleates local protein gradients to propagate silencing across the X chromosome. Cell. 184, 6174–6192.e32 (2021).

32. A. Cerase, A. Armaos, C. Neumayer, P. Avner, M. Guttman, G. G. Tartaglia, Phase separation drives X-chromosome inactivation: a hypothesis. Nat Struct Mol Biol. 26, 331–334 (2019).

33. A. Pandya-Jones, Y. Markaki, J. Serizay, T. Chitiashvili, W. R. Mancia Leon, A. Damianov, C. Chronis, B. Papp, C. K. Chen, R. McKee, X. J. Wang, A. Chau, S. Sabri, H. Leonhardt, S. Zheng, M. Guttman, D. L. Black, K. Plath, A protein assembly mediates Xist localization and gene silencing. Nature. 587, 145–151 (2020).

34. J. W. Jachowicz, M. Strehle, A. K. Banerjee, M. R. Blanco, J. Thai, M. Guttman, Xist spatially amplifies SHARP/SPEN recruitment to balance chromosome-wide silencing and specificity to the X chromosome. Nat Struct Mol Biol. 29, 239–249 (2022).

35. B. Mészáros, G. Erdös, Z. Dosztányi, IUPred2A: Context-dependent prediction of protein disorder as a function of redox state and protein binding. Nucleic Acids Res. 46, W329–W337 (2018).

36. A. S. Belmont, Nuclear Compartments: An Incomplete Primer to Nuclear Compartments, Bodies, and Genome Organization Relative to Nuclear Architecture. Cold Spring Harb Perspect Biol, a041268 (2021).

37. B. Zhao, A. Katuwawala, C. J. Oldfield, G. Hu, Z. Wu, V. N. Uversky, L. Kurgan, Intrinsic Disorder in Human RNA-Binding Proteins. J Mol Biol. 433, 167229 (2021).

38. V. N. Uversky, Recent Developments in the Field of Intrinsically Disordered Proteins: Intrinsic Disorder-Based Emergence in Cellular Biology in Light of the Physiological and Pathological Liquid-Liquid Phase Transitions. Annu Rev Biophys. 50, 135–156 (2021).

39. S. Al Nadaf, P. D. Waters, E. Koina, J. E. Deakin, K. S. Jordan, J. A. M. Graves, Activity map of the tammar X chromosome shows that marsupial X inactivation is incomplete and escape is stochastic. Genome Biol. 11, R122 (2010).

40. C. L. Rodríguez-Delgado, S. A. Waters, D. P. Waters, Paternal X inactivation does not correlate with X chromosome evolutionary strata in marsupials. BMC Evol Biol. 14, 4–11 (2014).

41. N. C. Lister, A. M. Milton, H. R. Patel, S. A. Waters, K. L. McIntyre, A. M. Livernois, L. Kian Wee, A. R. Ringel, S. Mundlos, M. I. Robson, L. Shearwin-Whyatt, F. Grützner, J. A. M. Marshall Graves, A. Ruiz-Herrera, P. D. Waters, Incomplete transcriptional dosage compensation of vertebrate sex chromosomes is balanced by post-transcriptional compensation. bioRxiv, 312–320 (2023).

42. G. Su, A. Kuchinsky, J. H. Morris, D. J. States, F. Meng, GLay: Community structure analysis of biological networks. Bioinformatics. 26, 3135–3137 (2010).

43. H. Peterson, L. Kolberg, U. Raudvere, I. Kuzmin, J. Vilo, gprofiler2 -- an R package for gene list functional enrichment analysis and namespace conversion toolset g: Profiler. F1000Res. 9, 1–27 (2020).

